# The GTPase Nog1 couples polypeptide exit tunnel quality control with ribosomal stalk assembly

**DOI:** 10.1101/462333

**Authors:** Purnima Klingauf-Nerurkar, Ludovic Gillet, Cohue Peña, Olga T. Schubert, Martin Altvater, Yiming Chang, Ruedi Aebersold, Vikram Govind Panse

**Affiliations:** Institute of Medical Microbiology, University of Zurich, CH-8006 Zurich, Switzerland; Institute of Molecular Systems Biology, ETH Zurich, CH-8093 Zurich, Switzerland; Current address: Department of Human Genetics, University of California, Los Angeles, Los Angeles, CA 90095, USA; Institute of Biochemistry, ETH Zurich, CH-8093 Zurich, Switzerland; Faculty of Science, University of Zurich, CH-8057, Switzerland

## Abstract

Eukaryotic ribosome precursors acquire translation competence in the cytoplasm through stepwise release of bound assembly factors, and proofreading of their functional centers. In case of the large subunit precursor (pre-60S), these essential steps include eviction of placeholders Arx1 and Mrt4 that prevent premature loading of the protein-folding machinery at the polypeptide exit tunnel (PET), and the ribosomal stalk, respectively. Here, we reveal that sequential ATPase and GTPase activities license release factors Rei1 and Yvh1 recruitment to the pre-60S in order to trigger Arx1 and Mrt4 removal. Drg1-ATPase activity extracts the C-terminal tail of Nog1 from the PET, enabling Rei1 to probe PET integrity, and then catalyze Arx1 release. Subsequently, GTPase hydrolysis stimulates Nog1 removal from the pre-60S, permitting Yvh1 to mediate Mrt4 release, and initiate ribosomal stalk assembly. Thus, Nog1 couples quality control and assembly of spatially distant functional centers during ribosome formation.

## Introduction

Error-free translation of the genetic code by the ribosome is critical for proteome homeostasis and cellular function. This essential task necessitates that only correctly assembled ribosomal subunits are committed for translation.

The eukaryotic ribosome is made up of a large 60S subunit containing 25S, 5.8S and 5S ribosomal RNAs (rRNAs) and 46 ribosomal proteins (r-proteins), and a small 40S subunit containing 18S rRNA and 33 r-proteins (Ben-Shem et al., 2011). Eukaryotic ribosome assembly is initiated by RNA-polymerase I driven production of pre-rRNA in the nucleolus (Kressler et al., 2017; Pena et al., 2017). The emerging pre-rRNA associates with 40S specific r-proteins, assembly factors (UTPs), U3 snoRNP and snoRNAs to form a pre-40S particle. Endonucleolytic cleavage releases the earliest pre-40S particle, permitting the remaining growing pre-rRNA to recruit 60S specific r-proteins and assembly factors to form a pre-60S particle. On their way through the nucleoplasm, pre-ribosomal particles interact with >200 assembly factors including >40 energy-consuming AAA-ATPases, ABC-ATPases, GTPases and ATP-dependent RNA helicases. Elucidating their order of action, and the mechanisms that co-ordinate their activities during pre-ribosome maturation is an important challenge.

Export-competent pre-ribosomes are transported through nuclear pore complexes (NPCs) into the cytoplasm where they undergo maturation and proofreading before acquiring translation competence (Nerurkar et al., 2015). These steps include release of assembly factors, transport receptors, pre-rRNA processing steps and incorporation of remaining r-proteins critical for ribosome function. Cytoplasmic maturation of a pre-60S particle is initiated by the AAA-ATPase Drg1, which evicts the placeholder ribosomal-like protein Rlp24 from the pre-60S particle (Kappel et al., 2012; Kressler et al., 2012; Pertschy et al., 2007). This is a prerequisite for subsequent release of GTPases Nog1 and Nug1, the assembly factor Nsa2, the 40S anti-association factor Tif6, the ribosomal-like protein Mrt4 and export factors Mex67-Mtr2, Bud20, Arx1 and Nmd3 (Loibl et al., 2014; Zisser et al., 2018). Following Rlp24 release, the 60S maturation pathway, *via* a yet unknown mechanism, bifurcates to proofread the polypeptide exit tunnel (PET) and assemble the ribosomal stalk (Lo et al., 2010). PET maturation is accomplished through the release of Arx1, an assembly factor that seals the ribosomal tunnel, and prevents premature loading of the protein-folding chaperone machinery on immature pre-60S particles (Bradatsch et al., 2012; Greber et al., 2012). Arx1 release from the pre-60S particle requires the cytoplasmic zinc-finger protein Rei1, the DnaJ domain-containing protein Jjj1and Ssa1/Ssa2 (Hsp70) ATPase activity (Meyer et al., 2010; Meyer et al., 2007). Cryo-electron microscopy (cryo-EM) studies revealed that Rei1 deeply inserts its C-terminal tail (Rei1-CTT) into the PET (Greber et al., 2016). Failure to insert Rei1-CTT into the tunnel impairs Arx1 release, and blocks further pre-60S particle maturation (Greber et al., 2016). Curiously, cryo-EM analyses of a nuclear pre-60S particle revealed that the GTPase Nog1 inserts it’s C-terminal tail (Nog1-CTT) into the ribosomal tunnel during nuclear biogenesis steps (Wu et al., 2016). For Rei1 to gain access to the ribosomal tunnel, the Nog1-CTT must be extracted. However, it is unclear how these events are coordinated during final cytoplasmic maturation.

Another functional center whose assembly is completed in the cytoplasm is the 60S ribosomal stalk. The stalk is built from a single copy of ribosomal proteins uL10 (Rpp0) and two heterodimers of P1 (Rpp1) and P2 (Rpp2), and plays an essential role during translation by recruiting and activating translation elongation factors. On a mature 60S subunit, the stalk is anchored through the interaction of uL10 with rRNA and uL11 (Rpl12) (Ben-Shem et al., 2011). During 60S assembly, the ribosomal-like protein Mrt4 functions as a placeholder for uL10 (Kemmler et al., 2009; Lo et al., 2009). Mrt4 removal from the pre-60S is triggered by the release factor Yvh1 *via* a yet unknown mechanism. Only after Mrt4 is released, uL10 can be loaded onto the pre-60S, to assemble the ribosomal stalk.

Removal of placeholders Arx1 and Mrt4 from the pre-60S particle is essential for progression of the cytoplasmic maturation pathway. These events are critical to acquire translation competence through the release of anti-40S subunit joining factors Tif6 and Nmd3. While Drg1 AAA-ATPase activity is essential for both PET maturation/quality control and stalk assembly, it is unclear how these events located on the opposite ends of pre-ribosome are orchestrated. By employing a combination of state of the art quantitative mass spectrometry, genetic and cell-biological approaches, we reveal that sequential Drg1-ATPase and Nog1-GTPase activities license Arx1 and Mrt4 removal from the pre-60S particle. Drg1 extracts the Nog1-CTT from the ribosomal tunnel, permitting Rei1 to probe its integrity, a key step to trigger Arx1 release. Nog1 then stimulates its own removal from the pre-60S particle permitting Yvh1 to mediate Mrt4 release and initiate ribosomal stalk biogenesis. Thus, Nog1 couples quality control and assembly of spatially distant functional centers during 60S formation.

## Results

The nuclear-localized GTPase Nog1 co-enriches with late pre-60S particles that contain export factors Nmd3, Bud20 and Arx1 (Jensen et al., 2003; Kallstrom et al., 2003). In WT cells, Nog1 is not detected on a cytoplasmic particle purified through Tandem-affinity purification (TAP) of Lsg1 (Figure S1; Altvater et al., 2012; Kressler et al., 2008). However, it mis-localizes to the cytoplasm, and accumulates on Lsg1-TAP particles upon impairment of the cytoplasmic AAA-ATPase Drg1 or upon treating yeast cells with the Drg1-inhibitor diazaborine (DIA) (Loibl et al., 2014; Pertschy et al., 2007). These data indicate that Nog1 travels to the cytoplasm, where it is released in a Drg1-dependent manner and rapidly recycled back to the nucleus for another biogenesis round. How, and exactly when Nog1 is released from the pre-60S particle in the cytoplasm is unclear.

### Nog1^DN^ accumulates on a cytoplasmic pre-60S particle

The Nog1 G-domain exhibits characteristic G1-G5 motifs (Figure 1A) suggesting that like other pre-60S associated GTPases such as Nug2 and Lsg1, GTP-binding or hydrolysis might regulate interactions between Nog1 and the pre-60S particle (Hedges et al., 2005; Matsuo et al., 2014). Dominant-negative mutations have been described within the G1 motif of Lsg1 (K349N/R/T) (Hedges et al., 2005) and in the G3 motif of Nug2 (G369A) (Bourne et al., 1991; Hedges et al., 2005; Matsuo et al., 2014) that impair the release of these GTPases from pre-60S particles. A G224A mutation in the G3 motif of human Nog1, which presumably blocks GTP hydrolysis, was shown to be dominant negative in mammalian cells, and induced pre-rRNA processing and assembly defects (Lapik et al., 2007).

**Figure 1.**
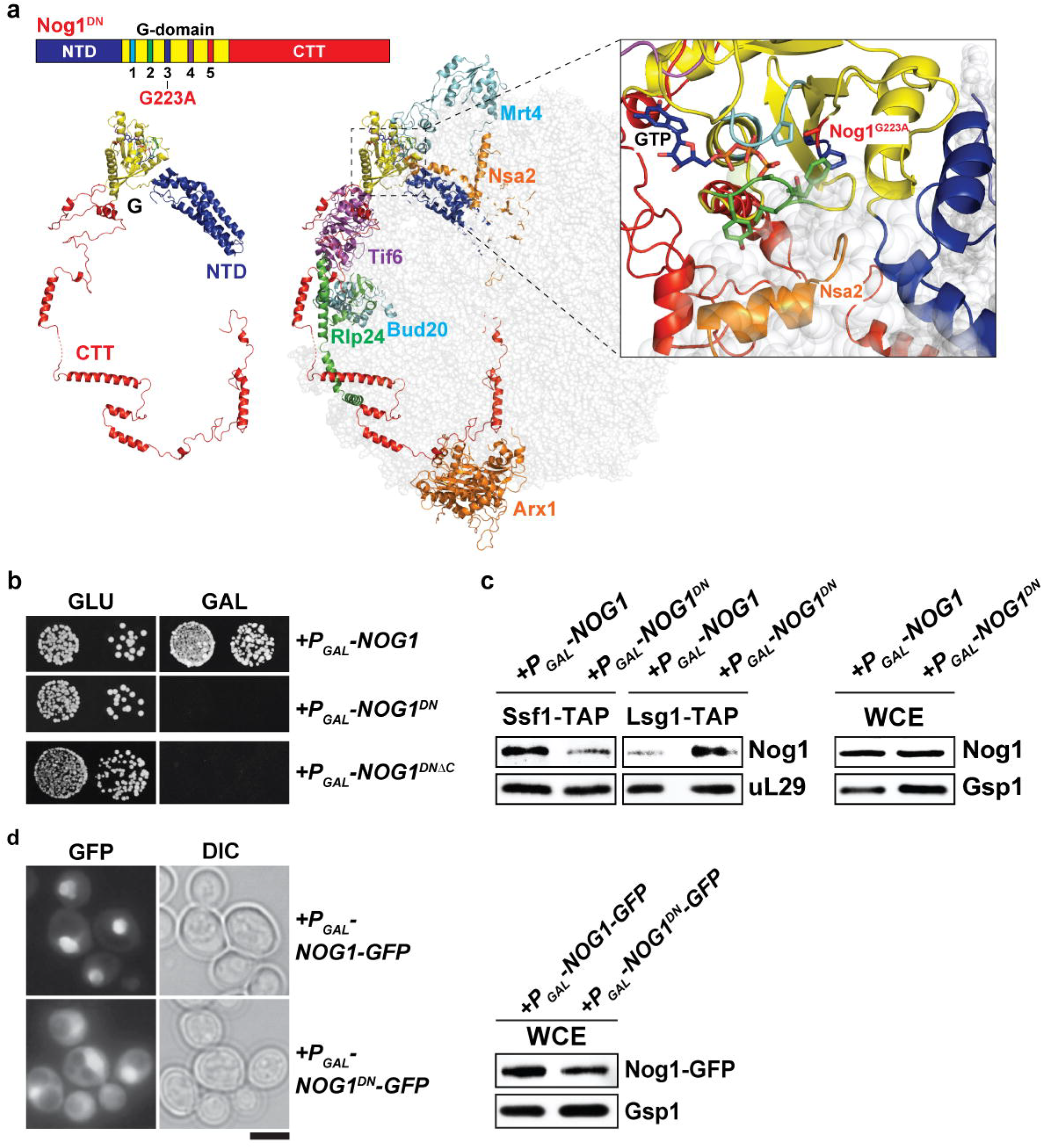
*Nog1^DN^* accumulates on a cytoplasmic pre-60S particle. **(A)** Nog1 domain organization and cryo-EM structure of a Nog1-containing pre-60S ribosome (PDB-3JCT). Selected assembly factors bound to the pre-60S are shown in color (Mrt4, Nsa2, Tif6, Rlp24, Bud20, Arx1). The Nog1G223A mutation is located in the G3 motif of its GTPase domain. **(B)** *NOG1-G223A* is dominant negative. BY wild-type cells were transformed with a plasmid encoding *NOG1* or *NOG1-G223A* under control of the galactose-inducible GAL1 promoter and spotted in 10-fold dilutions on glucose- or galactose-containing media. Plates were incubated at 30°C for 2-4 days. Nog1^DNΔC^ was truncated after residue D479 **(C)** Nog1-G223A accumulates on cytoplasmic pre-60S particles. Early nuclear (Ssf1-TAP) and cytoplasmic (Lsg1-TAP) pre-60S particles were purified from Nog1 or Nog1-G223A-expressing cells and analyzed by western blotting using indicated antibodies. Whole-cell extracts (WCE) served as control for Nog1 expression. **(D)** Nog1G223A mis-localizes to the cytoplasm. BY wild-type cells expressing Nog1-GFP or Nog1-G223A-GFP under control of GAL1 were grown in raffinose-containing synthetic medium to early log phase and then supplemented with 2% galactose. After 30min incubation, cells were washed, incubated in YPD for 3h and then visualized by fluorescence microscopy. Scale bar = 5 μm. Whole-cell extracts (WCE) served control for Nog1 expression.

We investigated whether the G-domain contributes to Nog1 release from the pre-60S particle in the cytoplasm. To this end, the orthologous mutation in the G3 motif (G223A) of yeast Nog1, hereafter termed Nog1^DN^ was transformed into a Nog1 shuffle strain, wherein the *NOG1* gene was disrupted, but viability of the yeast cells maintained through a centromeric plasmid containing a wild-type (WT) copy of *NOG1*. We did not obtain transformants for the Nog1^DN^ mutant. To demonstrate dominant negative behavior, we placed the Nog1^DN^ mutant under control of an inducible *GAL1* promoter, and transformed this plasmid into WT yeast cells. On glucose-containing medium, where Nog1^DN^ expression is repressed, the resulting transformants grew similar to WT. In contrast, expression of Nog1^DN^ in galactose-containing medium was lethal to yeast cells (Figure 1B), confirming the dominant negative behavior of the G223A mutation. Notably, expression of Nog1^DN^ lacking the C-terminal tail (CTT) (Nog1^DNΔC^) was also lethal in yeast (Figure 1B), implicating an impaired G3-domain for the dominant negative phenotype.

Nog1 is recruited to the pre-60S particle in the nucleolus (Altvater et al., 2012; Kressler et al., 2008), and is released from the particle in the cytoplasm (Altvater et al., 2012; Lo et al., 2010; Pertschy et al., 2007). We investigated whether the Nog1^DN^ mutant was efficiently released from the pre-60S in the cytoplasm. We isolated the Lsg1-TAP particle after inducing expression of either WT Nog1 or the Nog1^DN^ mutant allele for 2.5 hours (Figure 1C). Western analyses revealed that Nog1^DN^ mutant protein, but not WT Nog1 accumulated on the Lsg1-TAP particle (Figure 1C). Corresponding whole cell extracts (WCE) revealed similar WT Nog1 and Nog1^DN^ protein levels (Figure 1C), suggesting Nog1^DN^ co-enrichment with Lsg1-TAP is not due to altered expression of the mutant protein. The Nog1^DN^-GFP fusion also showed a significant increase in cytoplasmic signal supporting the notion that release of the mutant protein from the pre-60S in the cytoplasm is impaired (Figure 1D). Although a nuclear signal of Nog1^DN^-GFP is observed in these cells, this mutant did not efficiently co-enrich with Ssf1-TAP under the same conditions, possibly due to blockage in earlier pre-60S assembly steps (see later).

Based on all the above data, we conclude that an impaired G3 motif impairs Nog1 release from the pre-60S particle in the cytoplasm.

### Nog1^DN^ impairs cytoplasmic maturation of the pre-60S particle

We investigated consequences of impaired Nog1^DN^ release on the composition of the Lsg1-TAP particle. For this, we employed <u>S</u>equential <u>W</u>indow <u>A</u>cquisition of all <u>TH</u>eoretical fragment ion spectra mass spectrometry, also termed SWATH-MS. SWATH-MS is a mass spectrometry (MS) approach that combines data-independent acquisition with a peptide-centric data query strategy (Gillet et al., 2012). In contrast to selected reaction monitoring mass spectrometry (SRM-MS) (Picotti and Aebersold, 2012), SWATH-MS can be extended to the analysis of any peptide and protein of interest post-acquisition, while maintaining optimal consistency of quantification in pull-down samples (Collins et al., 2013; Lambert et al., 2013). We first interrogated quantitatively the protein composition of four well-characterized pre-60S particles at different maturation stages (Nissan et al., 2002): Ssf1-TAP, an early nucleolar particle; Rix1-TAP, a nucleoplasmic particle; Arx1-TAP, an export-competent particle and Lsg1-TAP, an exclusively cytoplasmic particle. The data was analyzed using OpenSWATH software (Rost et al., 2014), and accuracy was compared with previous SRM-MS based analyses (Altvater et al., 2012). We found that the proteomic heat map obtained *via* SWATH-MS was in excellent agreement with SRM-MS (Altvater et al., 2012) and Western analyses (Figure S1). In contrast to SRM-MS, SWATH-MS permitted quantitation and clustering of nearly all assembly factors that are known to interact with pre-60S particles (Figure 2).

**Figure 2.**
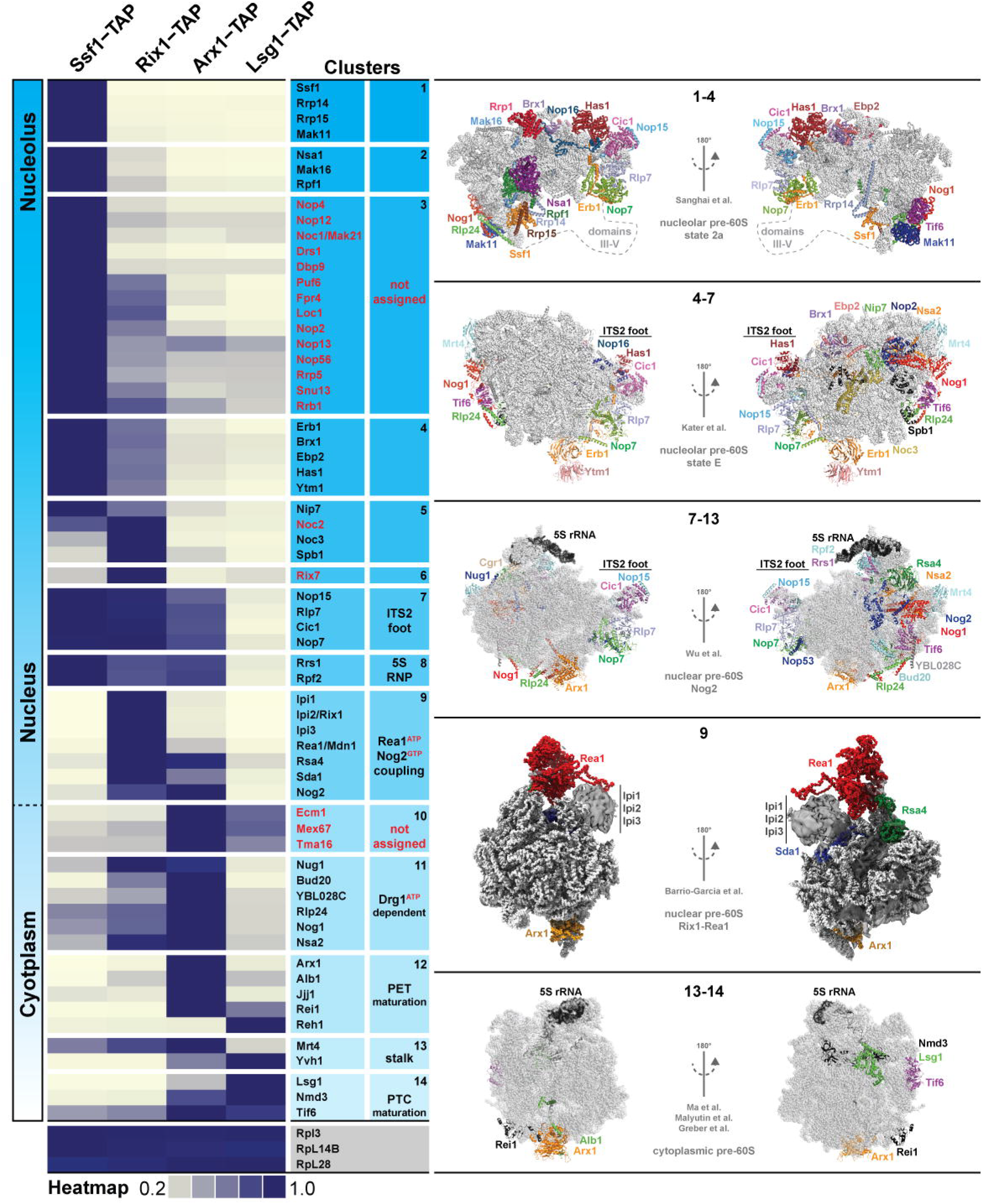
Dynamic association of assembly factors with pre-60S subunits as revealed by SWATH-MS. Purified pre-60S particles representing nucleolar (Ssf1-TAP), nuclear (Rix1-TAP), nuclear to cytoplasmic (Arx1-TAP) and cytoplasmic (Lsg1-TAP) stages were analyzed by SWATH-MS. The heatmap was generated based on the average of three independent biological replicates and depicts the relative and individual enrichment of 63 assembly factors with maturing pre-60S particles. The acquired SWATH data was normalized based on the average intensities of three depicted 60S r-proteins and the intensity of each factor was scaled to the highest intensity of that factor in the selected condition represented in the heatmap (Altvater et al.) Maximum enrichment is depicted as purple and minimum enrichment as gold. Assembly factors were then manually grouped into structure-functional clusters based on available structural information (nucleolar pre-60S: PDB-6C0F and PDB-6ELZ; nuclear pre-60S: PDB-3JCT and PDB-5FL8; cytoplasmic pre-60S: PDB-5H4P, PDB-5T62 and PDB-5APN). Factors currently not assigned in available pre-60S structures are marked red.

Next, we correlated the protein heat map with reported pre-ribosome cryo-EM structures at different assembly stages (Barrio-Garcia et al., 2016; Greber et al., 2016; Kater et al., 2017; Ma et al., 2017; Malyutin et al., 2017; Sanghai et al., 2018; Wu et al., 2016). These analyses allowed classification and organization of assembly factors into different clusters, and determination of their approximate residence time on the maturing pre-60S particle. For e.g. the early nucleolar Ssf1-Rrp14-Rrp15-Mak11 cluster, the nuclear ITS2 factors Nop15-Rlp7-Cic1-Nop7, the 5S RNP-associated Rrs1-Rpf2, the Rix1-Rea1 machinery and the Drg1-dependent factors Rlp24-Nog1-Nsa2 appear to undergo coordinated and grouped release. In addition, our analyses revealed temporal association of numerous assembly factors for which structural information in the context of the pre-60S is currently lacking. These factors include several nucleolar proteins (Figure 2, marked in red), but also a previously uncharacterized nuclear localized Tma16 (Figure 2). Altogether, we conclude that SWATH-MS is reliable tool to quantify the protein contents of pre-ribosomal particles.

Next, we quantified and compared the protein contents of two genetically trapped cytoplasmic pre-60S particles isolated from yeast cells expressing either the dominant negative Drg1-E617Q, (Lsg1-TAP:Drg1^DN^) (Altvater et al., 2012) or Nog1^DN^ (Lsg1-TAP:Nog1^DN^) mutant. We focused on assembly factors known to participate in pre-60S cytoplasmic maturation (Figure 3A). In agreement with previous studies (Altvater et al., 2012; Lo et al., 2010; Pertschy et al., 2007), a Drg1^DN^:Lsg1-TAP particle accumulated Rlp24, Bud20, Nog1, Mrt4, Nmd3 and Tif6. In contrast, we found that the Lsg1-TAP:Nog1^DN^ particle accumulated Mrt4, Nmd3 and Tif6, but not Rlp24 and Bud20. Both Drg1^DN^ and Nog1^DN^-trapped Lsg1-TAP particles failed to recruit Yvh1. Intriguingly, while Rei1 recruitment to Lsg1-TAP was impaired upon Drg1^DN^ expression, Nog1^DN^-trapped particles, in contrast, accumulated Rei1. The relative co-enrichments of assembly factors on the Lsg1-TAP:Nog1^DN^ particle monitored *via* SWATH-MS were in agreement with Western analyses (Figure 3C). These biochemical data suggest that Nog1^DN^ expression impairs a subset of events along the 60S cytoplasmic maturation pathway.

**Figure 3.**
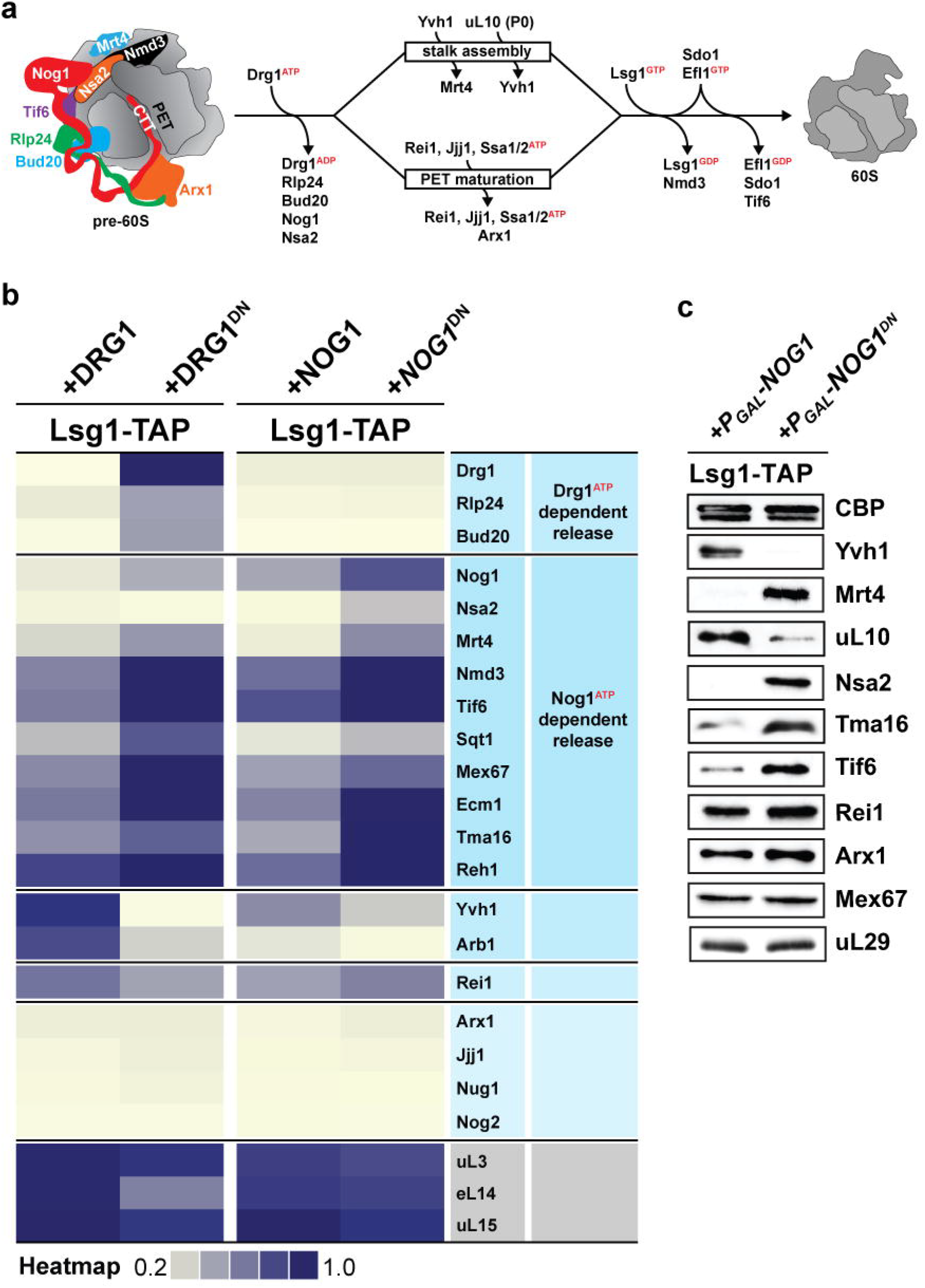
*Nog1^DN^* impairs cytoplasmic maturation of the pre-60S particle. **(A)** Current model for cytoplasmic maturation of the pre-60S subunit. A cytoplasmic pre-60S is shown as cartoon representation based on Figure 1a. Energy-consuming assembly factors are indicated with ATP or GTP. **(B)** SWATH-MS analysis of cytoplasmic Lsg1-TAP pre-60S particles upon expression of *DRG1* or *NOG1* dominant negative alleles. Assembly factors accumulating on the pre-60S particle in a Drg1- or Nog1-dependent manner were clustered based on increased relative enrichment in trapped particles. Analysis was performed as in Figure 2. The intensity of each factor was scaled to the highest density of that factor in the four previous purifications of Figure 2. **(c)** Western analyses of selected pre-60S assembly factors. *Nog1^DN^*-trapped Lsg1-TAP was purified as in (b) and then subjected to western analysis using indicated antibodies. CBP: calmodulin binding peptide present in the Lsg1 TAP-tag. Asterisk: unspecific background band.

### Nog1^DN^ does not hinder initiation of cytoplasmic maturation

To dissect the impact of Nog1^DN^ accumulation on a pre-60S particle during each step along the cytoplasmic maturation pathway, we employed a cell-biological approach. Following nuclear export, the C-terminal region within Rlp24 recruits Drg1 and stimulates its ATPase activity, resulting in Rlp24 release from the pre-60S particle (Altvater et al., 2012; Kappel et al., 2012; Lo et al., 2010; Pertschy et al., 2007) (Figure 3A). Since Rlp24 intertwines with the Nog1-CTT (Wu et al., 2016) (Figure 1A), we wondered whether Nog1^DN^ hinders Rlp24 release from a pre-60S particle. To test this, we monitored Rlp24-TAP localization by immunofluorescence after induction of Nog1^DN^ expression. Rlp24-TAP did not mis-localize to the cytoplasm upon Nog1^DN^ induction (Figure 4A), suggesting that Nog1^DN^ accumulation on the cytoplasmic pre-60S particle did not affect the release of Rlp24 and recycling back to the nucleus. These data are consistent with SWATH-MS and Western analyses which show that Rlp24 does not accumulate on the Lsg1-TAP:Nog1^DN^ particle (Figure 3B). We conclude that expression of Nog1^DN^ does not disturb initiation of the cytoplasmic maturation cascade. Our finding is in agreement with previous studies demonstrating that Drg1-ATPase activity is necessary and sufficient to release Rlp24 from the pre-60S particle (Kappel et al., 2012). Consistent with SWATH-MS, Bud20 and Nug1 release from the pre-60S in the cytoplasm were also not impaired upon Nog1^DN^ expression (Figure 4B).

**Figure 4.**
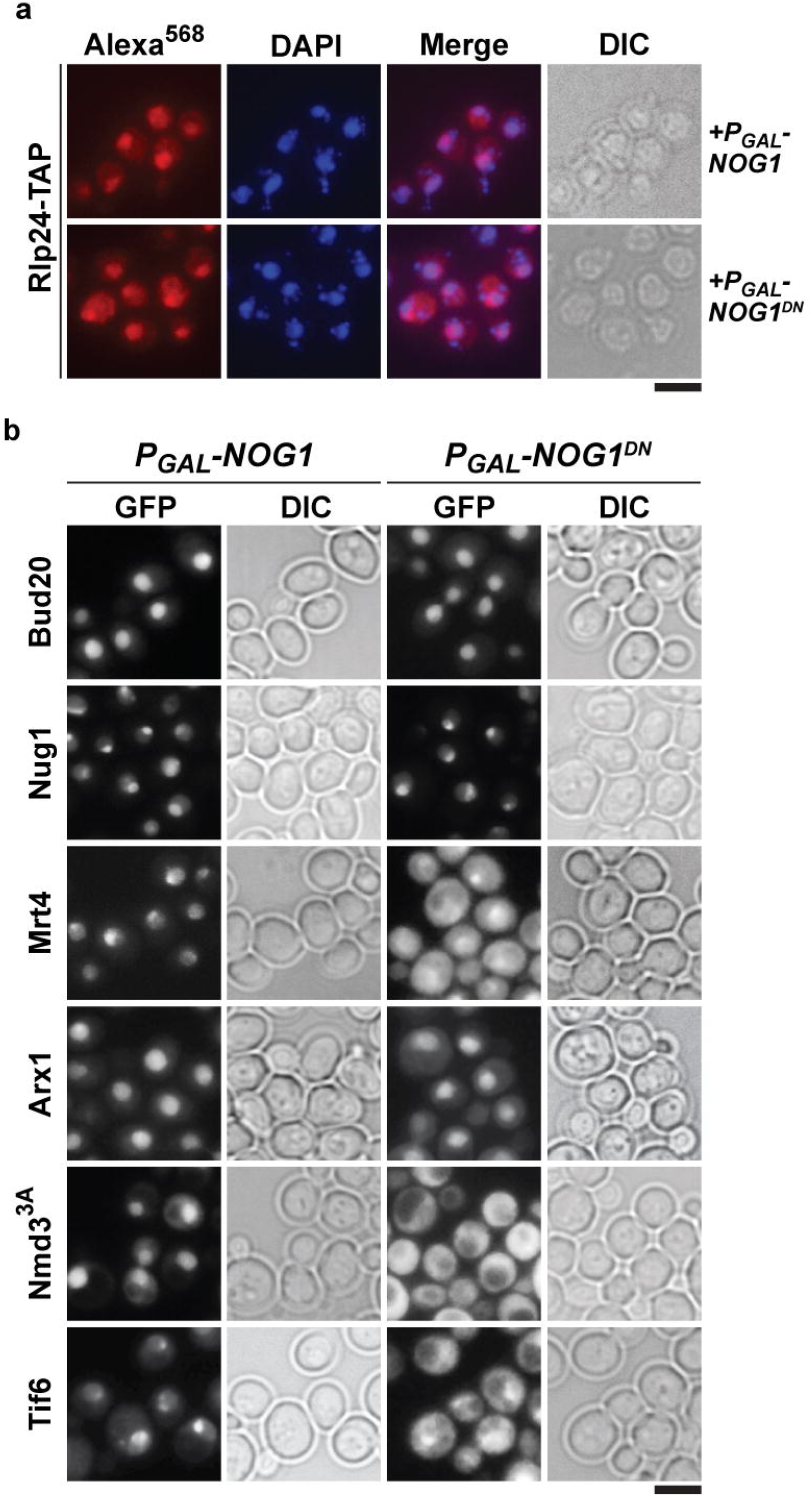
*Nog1^DN^* does not hinder initiation of cytoplasmic maturation. Mrt4, Nmd3^3A^ and Tif6 mis-localize to the cytoplasm upon expression of Nog1^DN^, but not Bud20, Nug1 and Arx1 **(A)** A *RLP24-TAP* strain expressing *NOG1* or *NOG1^DN^* under control of GAL1 was grown in raffinose-containing synthetic medium to early log phase, then supplemented with 2% galactose, and further grown for 3h. Cells were then fixed and prepared for indirect immunofluorescence using an anti-Protein A antibody. Cells were visualized by fluorescence microscopy. Scale bar = 5 μm. **(B)** Indicated GFP-tagged strains expressing *NOG1* or *NOG1^DN^* under control of GAL1 were grown as in (a) and then visualized by fluorescence microscopy. Scale bar = 5 μm.

While the Lsg1-TAP:Drg1^DN^ particle accumulates both Drg1^DN^ and Nog1, the Lsg1-TAP:Nog1^DN^ particle accumulates only Nog1^DN,^, but not Drg1 (Figure 3B). We infer that complete Nog1 release from a pre-60S particle occurs only after Drg1-dependent Rlp24 release, and requires a functional G3 motif.

### Nog1^DN^ interferes with ribosomal stalk assembly

After Rlp24 release, the pre-60S cytoplasmic maturation pathway bifurcates into two parallel pathways (Lo et al., 2010): (1) to assemble the ribosomal stalk and (2) to mature and proofread PET integrity (Figure 3A). Ribosomal stalk assembly occurs in the cytoplasm from uL10 (P0) and P1/P2 protein heterodimers (Gonzalo and Reboud, 2003). Mrt4 functions as a nuclear placeholder for uL10 (Rodriguez-Mateos et al., 2009) and joins the pre-60S already in the nucleolus (Lo et al., 2009). Mrt4 release in the cytoplasm requires Yvh1, which is maximally enriched on Lsg1-TAP (Kemmler et al., 2009). SWATH-MS and Western analyses revealed that Lsg1-TAP:Nog1^DN^ failed to recruit Yvh1, thereby inducing Mrt4 accumulation on cytoplasmic pre-ribosomes (Figure 3B, 3C). Consequently, recruitment of uL10 to Lsg1-TAP:Nog1^DN^ was also impaired (Figure 3C). In agreement with these data Nog1^DN^ expression mis-localises Mrt4-GFP to the cytoplasm (Figure 4B). Thus, Nog1^DN^ accumulation on the pre-60S particle prevents Yvh1 recruitment, a critical step to release Mrt4 and initiate ribosome stalk assembly. We conclude that ribosomal stalk assembly requires both Drg1-ATPase and Nog1-GTPase activities.

### Nog1^DN^ does not impair PET maturation and quality control

Next, we investigated the impact of Nog1^DN^ expression on the second branch of the cytoplasmic maturation pathway: PET maturation and quality control. During nuclear assembly the aminopeptidase fold of Arx1 functions as a placeholder to seal the PET (Bradatsch et al., 2012; Greber et al., 2012). Arx1 release is triggered in the cytoplasm by Rei1 (Meyer et al., 2010), which is recruited to the pre-60S particle following Drg1-mediated Rlp24 release (Altvater et al., 2012; Greber et al., 2016; Hung and Johnson, 2006). Similar to Nog1-CTT, Rei1 contains a C-terminal tail (Rei-CTT) that is inserted into the PET (Greber et al., 2016). Failure to insert Rei1-CTT into the PET for e.g. by attaching a bulky domain (*rei1-TAP* or *rei1-GFP*) to Rei1 blocks Arx1 release and progression of cytoplasmic maturation, suggesting that PET proofreading represents a quality control check point (Greber et al., 2016). Before Rei1-CTT can probe PET integrity, Nog1-CTT needs to be extracted from the PET in the cytoplasm. SWATH-MS and Western analyses of Lsg1-TAP:Nog1^DN^ revealed that Rei1 was recruited and even accumulates on this particle (Figure 3B,C). Also, Nog1^DN^ expression did not alter the localization of Arx1-GFP suggesting that it does not interfere with Rei1’s ability to probe the PET (Figure 4B). Consistent with this notion, Arx1-GFP mis-localized to the cytoplasm upon Nog1^DN^ expression in a *rei1-TAP* mutant (Figure 5A). These data show that presence of Nog1^DN^ on the pre-60S particle does not hinder Rei1 recruitment, and insertion of the Rei1-CTT into the PET.

**Figure 5.**
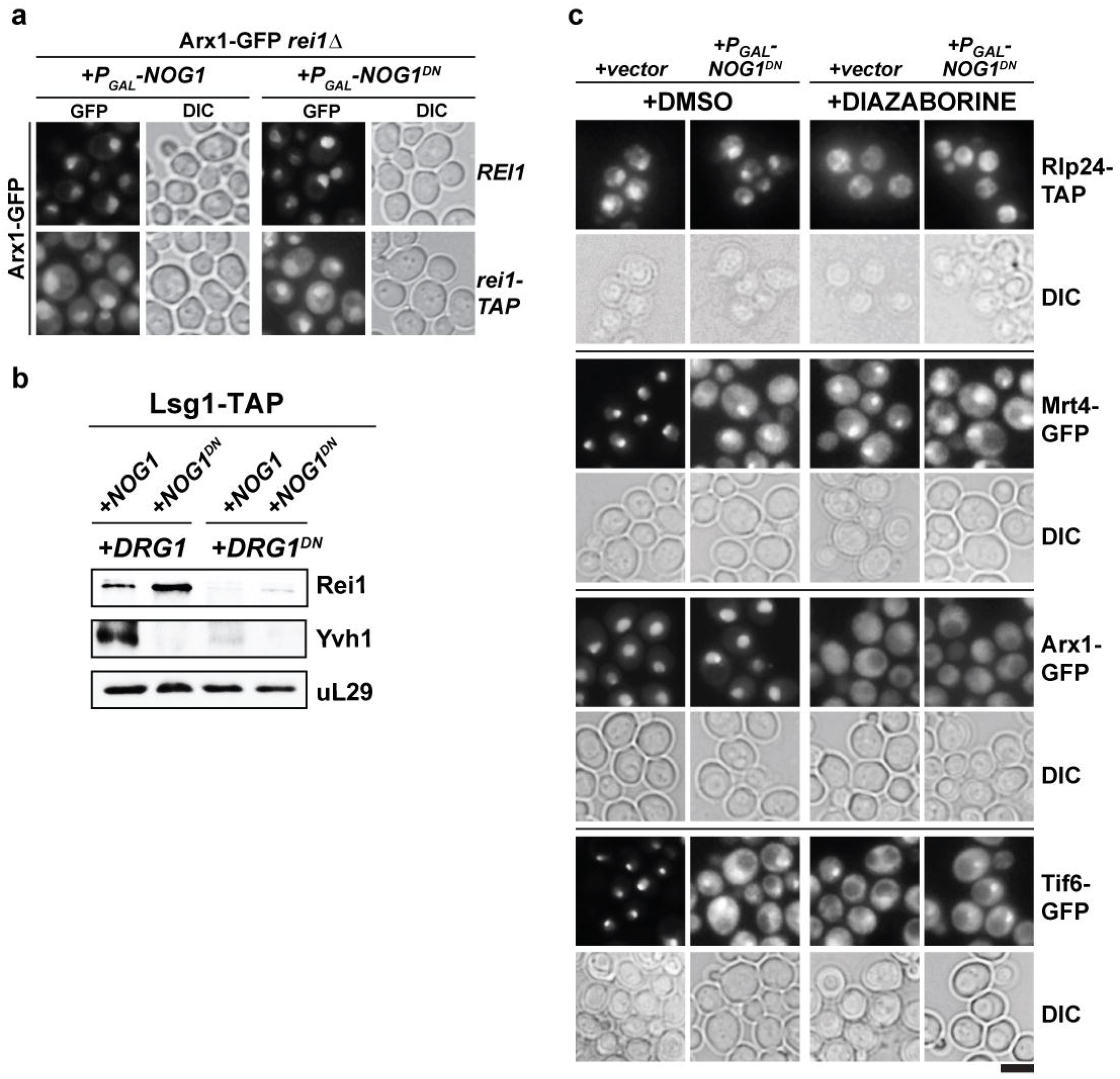
*NOG1^DN^* does not impair PET maturation and quality control. **(A)** Arx1 mis-localizes to the cytoplasm in *rei1Δ* or *rei1-TAP*, but not upon expression of *NOG1DN*. An *ARX1-GFP rei1Δ* strain was co-transformed with plasmids each expressing *REI1* or *rei1-TAP* under control of their native promoter and galactose-inducible *NOG1* or *NOG1^DN^*, respectively. Cells were grown in raffinose-containing synthetic medium to early log phase and then supplemented with 2% galactose. After 3h, cells were visualized by fluorescence microscopy. Scale bar = 5 μm. **(B)** *DRG1^DN^* impairs Yvh1 and Rei1 recruitment, whereas *NOG1^DN^* impairs Yvh1, but not Rei1 recruitment. A *LSG1-TAP* strain was co-transformed with plasmids each expressing copper-inducible *DRG1* or *DRG1^DN^* and galactose-inducible *NOG1* or *NOG1^DN^*, respectively. Cells were grown in raffinose-containing synthetic medium to early log phase and then supplemented with 0.5 mM copper sulfate and 2% galactose. After 3h, cells were lysed, purified through Lsg1-TAP and subjected to western analyses using indicated antibodies. **(c)** Indicated strains expressing *NOG1DN* under control of GAL1 were treated with DMSO or Diazaborine to block Drg1 activity and visualized by fluorescence microscopy as in Figure 4.

Since the Nog1-CTT and Rlp24 intimately intertwine around each other (Figure 1A), we investigated that Nog1-CTT extraction from the PET is linked to Drg1-mediated release of Rlp24 from the pre-60S. To this end, we monitored Rei1 recruitment to cytoplasmic pre-60S particles after co-expressing both Nog1^DN^ and Drg1^DN^. While Nog1^DN^ expression did not interfere with Rei1 recruitment, co-expression of both Nog1^DN^ and Drg1^DN^ impaired Rei1 recruitment to the pre-60S particle (Figure 5B). Similarly, Nog1^DN^ expression mis-localized Arx1-GFP to the cytoplasm in cells treated with the Drg1-inhibitor DIA, suggesting that Drg1-ATPase activity is critical for PET maturation/quality control, and precedes ribosomal stalk assembly (Figure 5C). As expected, expression of Drg1^DN^ alone or Nog1^DN^ alone impaired Yvh1 recruitment (Figure 5B). Further, Nog1^DN^ expression in DIA treated cells mis-localized Mrt4-GFP to the cytoplasm (Figure 5C). We conclude that Nog1^DN^ does not hinder cytoplasmic PET maturation and quality control.

### Nog1^DN^ blocks the terminal cytoplasmic maturation steps

The terminal steps during cytoplasmic maturation of a pre-60S particle include the coupled release of the nuclear export signal (NES) containing Crm1 adaptor Nmd3 and the 60S anti-association factor Tif6 (Weis et al., 2015). Nmd3 and Tif6 release from a pre-60S particle requires successful completion of PET maturation/quality control and ribosomal stalk assembly (Lo et al., 2010) (Figure 3A). Nog1^DN^ expression accumulated both Nmd3 and Tif6 on the pre-60S particle as judged by SWATH-MS and Western analyses (Figure 3B,C). To monitor Nmd3 accumulation on a cytoplasmic pre-60S particle by cell-biological means, we employed the nuclear localized Nmd3^3A^-GFP mutant fusion, which harbors mutations in its NES that decrease its nuclear export rate (Hedges et al., 2005). Upon induction of WT Nog1, as expected, both Nmd3^3A^-GFP and Tif6-GFP localize to the nucleus (Figure 4B). However, Nog1^DN^ expression mis-localized both factors to the cytoplasm (Figure 4B). We suggest that Nog1^DN^ expression impairs cytoplasmic release of Tif6 and Nmd3 from a pre-60S particle.

Previous studies showed that both Mrt4-GFP and Tif6-GFP mis-localize to the cytoplasm in *yvh1Δ* cells (Kemmler et al., 2009; Lo et al., 2009) (Figure 6A). A gain-of-function allele *mrt4^G68E^* that bypasses Yvh1-mediated Mrt4 release from the pre-60S particle rescued the slow growth of the *yvh1∆* mutant (Figure 6B,C), as well as Tif6-GFP localization (Figure 6A). Mrt4^G68^-GFP re-localized to the nucleus in the *yvh1∆* mutant (Kemmler et al., 2009) (Figure 6A). We investigated whether the *mrt4^G68E^* allele rescued the dominant negative behavior of Nog1^DN^. This was however not the case; Nog1^DN^ expression was dominant negative when expressed in *yvh1∆*Mrt4^G68E^ cells (Figure 6B). Further, Nog1^DN^ expression in *yvh1∆*Mrt4^G68E^ cells still mis-localized Tif6-GFP to the cytoplasm (Figure 6A). These results indicate that the presence of Mrt4 on cytoplasmic pre-60S particles is not the cause for Tif6-GFP mis-localization in Nog1^DN^ cells. Instead, the data implicate the presence of Nog1^DN^ on the pre-60S particle to block terminal steps leading to a translation competent 60S subunit.

**Figure 6.**
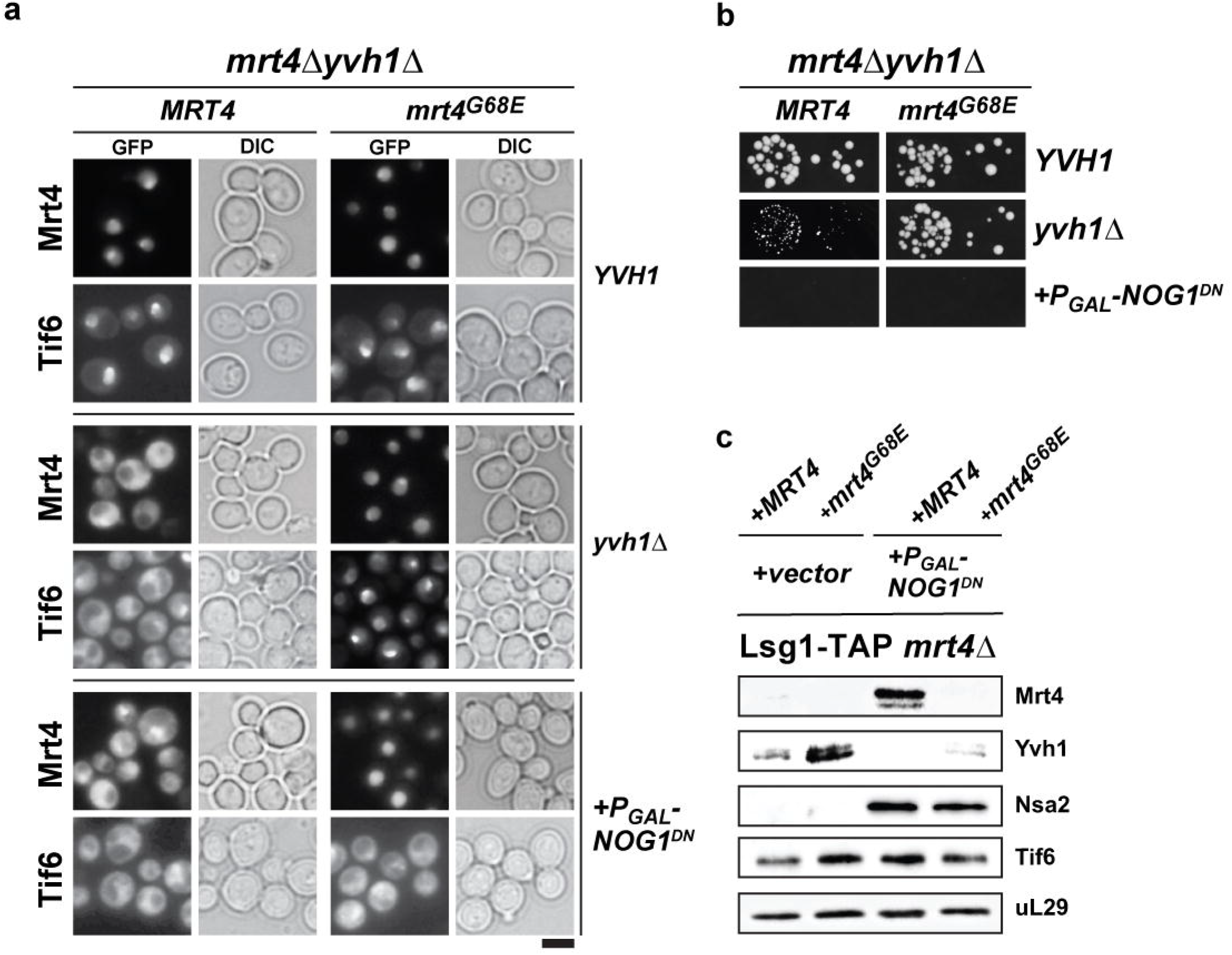
*NOG1^DN^* blocks the terminal cytoplasmic maturation steps. **(A)** *NOG1^DN^* impairs recycling of Mrt4 and Tif6. A *mrt4Δyvh1Δ* strain was co-transformed with plasmids encoding *MRT4-GFP* or *mrt4-G68E-GFP* and *YVH1* or galactose-inducible *NOG1^DN^*, respectively. To monitor Tif6 localization, the same strain was co-transformed with plasmids encoding TIF6-GFP, *MRT4* or *mrt4-G68E-GFP* and *YVH1* or galactose-inducible *NOG1^DN^*, respectively. Cells were grown in raffinose-containing synthetic medium to early log phase and then supplemented with 2% galactose. After 3h, cells were visualized by fluorescence microscopy. Scale bar = 5 μm. **(B)** The *mrt4-G68E* gain-of-function allele bypasses the need for *YVH1*, but does not rescue *NOG1^DN^*. The same strain as in (a) was spotted in 10-fold dilutions on galactose-containing media and incubated at 30°C for 2-4 days. **(C)** *NOG1^DN^* impairs recruitment of Yvh1, but not release of the Mrt4-G68E gain-of-function mutant. A *LSG1-TAP mrt4Δ* strain was co-transformed with plasmids encoding *MRT4* or *mrt4-G38E* and empty vector or galactose-inducible *NOG1^DN^*, respectively. Cells were grown in raffinose-containing synthetic medium to early log phase and then supplemented with 2% galactose. After 3h, cells were lysed, purified through Lsg1-TAP and subjected to western analyses using indicated antibodies.

## Discussion

During its journey from the nucleolus to the cytoplasm and its concomitant progression towards the mature ribosome, a pre-60S particle interacts with diverse essential energy-consuming enzymes. However, the precise order of recruitment/eviction and mechanisms that couple the different energy-consuming steps remain unknown. Here, we reveal an unanticipated functional coupling between an AAA-ATPase and a GTPase that ensures progression of a pre-60S towards translation competence.

Nog1 is an essential GTPase that travels into the cytoplasm with the pre-60S where it is released and recycled back into the nucleus (Jensen et al., 2003; Kallstrom et al., 2003). Our finding that the dominant negative G223A Nog1 mutant accumulates on Lsg1-TAP (Figure 1C) supports the notion that an impaired G-domain does not hinder recruitment to the assembling pre-60S particle. Instead, like other GTPases involved in other steps of 60S maturation (Hedges et al., 2005; Matsuo et al., 2014), impaired G3 domain blocks its release from the pre-60S particle. We and others (Lapik et al., 2007) have been unable to directly monitor GTPase activity of recombinant Nog1 *in vitro*. Similar to its bacterial homologue ObgE, it could be that GTP hydrolysis occurs only in context of the pre-ribosome (Feng et al., 2014). Nog1 is a predicted potassium-selective cation-dependent GTPase (also referred to as Group I CD-GTPase) (Ash et al., 2012). Group I CD-GTPases can be identified by the presence of two conserved asparagine residues in the G1 motif (Ash *et al.,* 2012). The first of the two conserved asparagine residues (Asn^K^) is coordinated to the potassium ion, whereas the second asparagine (Asn^SwI^) facilitates the Switch I structure. In yeast Nog1, Asn^K^ is present (N177), however, the second asparagine residue is substituted for an arginine (R185). This raises the possibility that the second asparagine (or equivalent) might be provided by a yet unknown activator protein. Based on cryo-EM studies (Wu et al., 2016), it is tempting to speculate that the neighboring assembly factor Nsa2 that makes intricate tertiary contacts with the GTPase domain might trigger GTPase activity upon undergoing a structural rearrangement (Figure 1A).

The equivalent G223A mutant in the G3 motif reported here is also dominant negative in the homologous Ras GTPase (Ford et al., 2005). Structural studies of this Ras mutant revealed that the switch region adopts an open conformation preventing GTP from inducing an active confirmation (Ford et al., 2005). Ras^G60A^ mutant was shown to sequester Ras GEFs into non-productive complexes leading to the depletion of the intracellular pool of available GEFs that is necessary to release GDP from Ras (Ford et al., 2005). It would be interesting to determine the molecular basis that underlies Nog1^DN^ dominant negative nature, and if it is similar to the Ras^G60A^ mutant.

Nog1, whose GTPase domain is docked at the P0 stalk base at a similar position as bacterial ObgE, is remarkably unique by its long C-terminal tail (CTT) which wraps around half the 60S subunit, passing by the peptidyl transferase center (PTC) and entering the polypeptide exit tunnel (PET) (Wu et al., 2016) (Figure 1A). Covering a total distance of >250 Å, the CTT interacts with several assembly factors Tif6, Rlp24, Bud20 and Arx1. Intriguingly, cryo-EM studies revealed that the Nog1-CTT intimately intertwines with the C-terminal alpha-helices of Rlp24 (Figure 1A). Despite this, we found that Rlp24 and Nog1 are independently released from the pre-60S particle, but follow a hierarchical order. Previous work showed that Drg1 grips Rlp24 *via* its C-terminus, and then triggers its ATP-dependent release from the pre-60S particle (Kappel et al., 2012). Drg1^DN^ expression *in vivo* prevents the release of Rlp24 and Nog1 from the pre-60S particle, suggesting that release of Nog1 requires prior Drg1 ATPase activity (Altvater et al., 2012; Pertschy et al., 2007). Based on its physical interactions with the Nog1-CTT, Rlp24 might serve an adaptor function for the AAA-ATPase Drg1 to extract Nog1-CTT from the PET.

Another factor that makes contacts with Rlp24 is the export factor Bud20 (Bassler et al., 2012). Like Rlp24, Drg1^DN^ expression interferes with Bud20 release and recycling from the pre-60S particle. It could be that release of Bud20 from the pre-60S particle is coupled and/or a consequence of Rlp24 release. Given that Bud20 localization is not altered upon Nog1^DN^ expression, we suggest that Bud20 release from the pre-60S particle occurs prior to bifurcation of the cytoplasmic maturation pathway.

Previous studies revealed that Drg1^DN^ expression impairs recruitment of Rei1 to the pre-60S particle and consequently Arx1 release (Altvater et al., 2012). In contrast, Nog1^DN^ expression does not interfere with PET maturation i.e. Rei1 recruitment to the pre-60S particle, proofreading of the ribosomal tunnel and Arx1 release. We infer that complete Nog1 eviction from the pre-60S particle is not required for this event. Instead, it appears that Rlp24-mediated extraction of the Nog1-CTT from the PET is sufficient to permit Rei1 recruitment, Rei1-CTT to probe PET integrity, and only then trigger Arx1 release.

While Nog1^DN^ expression *in vivo* did not interfere with PET maturation, we found that it impaired ribosomal stalk assembly. This assembly step requires prior release of Mrt4 that is facilitated by the release factor Yvh1 (Kemmler et al., 2009; Lo et al., 2009). We found that Nog1^DN^ expression interferes with Yvh1 recruitment to the pre-60S particle, and therefore impairing Mrt4 release. A cryo-EM structure of a cytoplasmic pre-60S particle revealed that Yvh1 docks between the central protuberance and stalk base adjacent to uL11 (Zhou et al. 2018). Together, these findings suggest that the presence of Nog1 and Nsa2 prevents the pre-60S from recruiting Yvh1, as both factors localize close to the stalk base and obstruct the Yvh1 RNA-binding site. These data also provide an explanation as to why failure to release Nog1 and Nsa2 from the pre-60S particle in the cytoplasm upon Drg1^DN^ expression also impairs ribosomal stalk assembly. While, Mrt4^G68E^ efficiently bypassed the need for Yvh1-mediated release, this gain-of-function allele did not rescue the dominant negative phenotype of Nog1^DN^. These cells were still impaired in the release of Tif6 from the pre-60S particle. We suggest that co-release of Nog1 and Nsa2 in the cytoplasm licenses a pre-60S particle to initiate ribosomal stalk assembly and downstream steps (Tif6 and Nmd3 release) essential to acquire translation competence.

Previously, the Johnson laboratory has ordered cytoplasmic maturation events on the pre-60S into a coherent pathway (Lo et al., 2010). In this pioneering study, the activity of AAA-ATPase Drg1 was proposed to initiate two parallel branches of the pathway i.e. PET maturation with quality control and ribosome stalk assembly. The two branches then converge to initiate Nmd3 and Tif6 release to acquire translational competence. Our data suggests that Nog1 plays a pivotal role in coordinating bifurcation of the 60S cytoplasmic maturation pathway. Recent cryo-EM structures and the data presented here permit refinement of the events that drive the cytoplasmic pre-60S maturation pathway (Figure 7). Upon arrival in the cytoplasm, the pre-60S recruits Drg1 via Rlp24’s C-terminal region. This interaction stimulates Drg1 ATPase activity resulting in Rlp24 removal, and consequently extraction of the Nog1-CTT from the PET. Removal of Rlp24 permits Rei1 recruitment via its N-terminal domain, whilst the Nog1 GTPase domain remains bound to the particle (Figure 7B). Following this event, Rei1-CTT probes PET integrity, and then triggers Arx1 release through the ATPases Ssa1/2 in order to complete tunnel maturation and quality control. Along the parallel pathway, extraction of the Nog1-CTT from the PET stimulates its GTP hydrolysis *via* a yet unknown mechanism, and therefore its complete release from the pre-60S particle. This step is crucial for Yvh1-mediated Mrt4 release and subsequent stalk assembly, but also allows the bifurcated pathway to converge and initiate terminal steps that drive Nmd3 and Tif6 release. While bound to the pre-60S, the N-terminal domain of Nog1 occupies the PTC region between Nmd3 and Tif6 and pushes 25S rRNA helix 89 outward (Figure 7C). This conformation clashes with the flexible N-terminal domain of Nmd3, which is prevented from moving across the PTC, and interact with Tif6, a crucial rearrangement step that initiates the coupled release of Nmd3 by Lsg1 and Tif6 by Efl1/Sdo1 (Ma et al., 2017; Malyutin et al., 2017; Weis et al., 2015; Yu et al.). Notably, recruitment of Lsg1 to the pre-60S is not dependent on initial cytoplasmic maturation steps, as we purified Drg1^DN^-trapped particles by Lsg1-TAP. Thus, Nog1 release from the pre-60S is essential for terminal steps to produce translation competent 60S subunits. We conclude that Nog1 serves as a hub that coordinates spatially distant maturation events to ensure correct 60S assembly through extensive contacts with multiple assembly factors.

**Figure 7.**
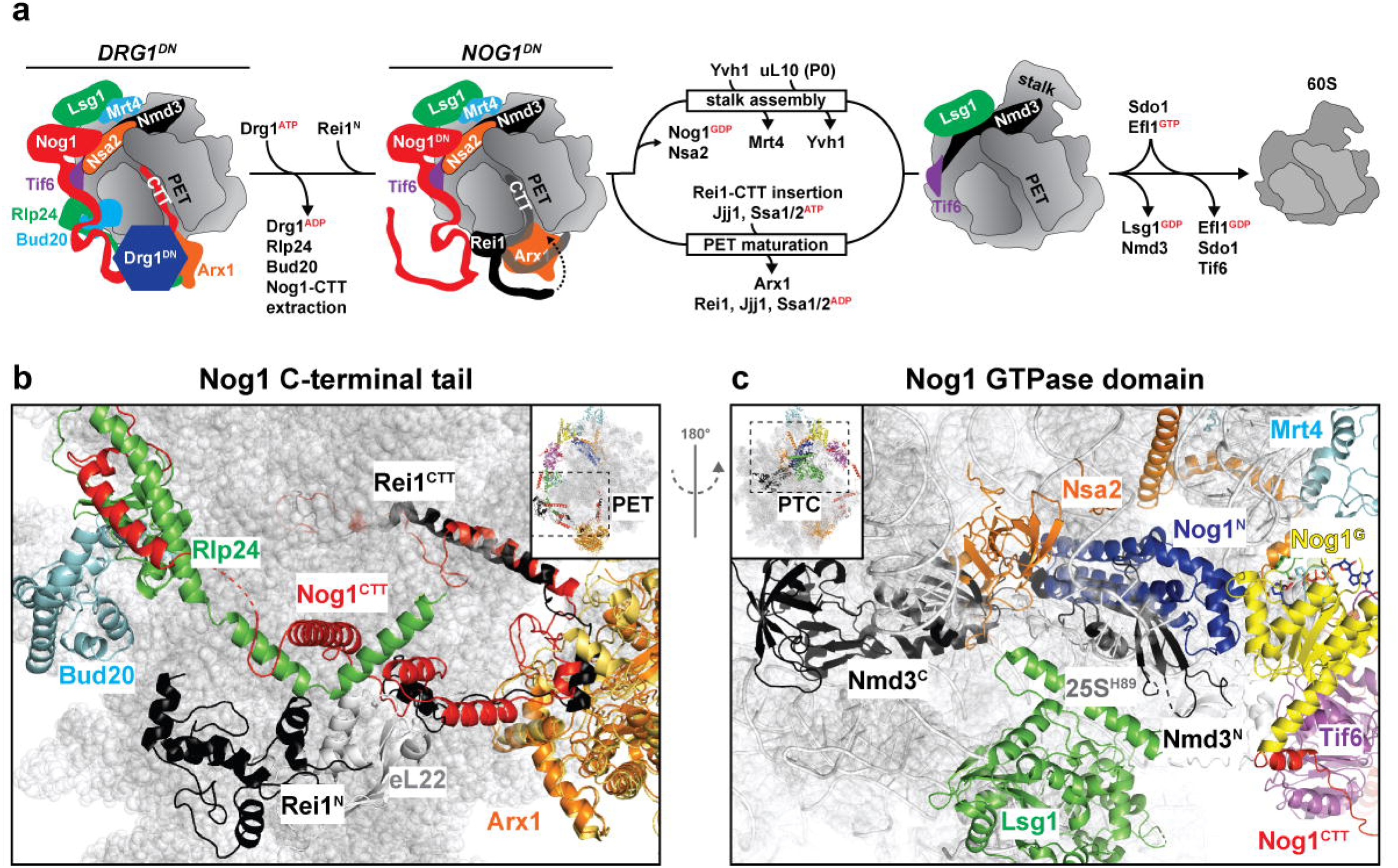
A revised model for the cytoplasmic maturation of the 60S pre-ribosome. **(A)** Schematic cartoon representation of the *DRG1^DN^*- and *NOG1^DN^*-trapped Lsg1-TAP particles analysed in this study. Drg1 initiates cytoplasmic maturation by releasing Rlp24 in an ATP-dependent manner. Removal of Rlp24 releases its interacting partner Bud20 and extracts the Nog1 C-terminal tail out of the polypeptide exit tunnel (PET). Nog1-CTT extraction allows, first, recruitment of the Rei1 N-terminal domain and, subsequently, insertion of the Rei1-CTT into the PET. The pathway then bifurcates into two independent events in which Rei1-CTT insertion triggers PET maturation through the ATP-dependent release of Arx1 by Ssa1/2 and Jjj1. Nog1-CTT extraction precedes the activation of Nog1 GTPase activity, which triggers its own release together with Nsa2. Release of these two factors allows recruitment of Yvh1 to release Mrt4 and initiate P0 stalk assembly. After PET maturation and stalk assembly, the pathway converges to perform the final steps to generate translation-competent 60S subunits. The remaining factors Nmd3 and Tif6 are each released by Lsg1 and Efl1/Sdo1 in a GTP-dependent manner, respectively. **(b)** The Rei1 N-terminus clashes with Rlp24 and the Rei1-CTT clashes with Nog1-CTT. Cryo-EM structures of nuclear Nog1-containing (PDB-3JCT) and late cytoplasmic Rei1-containing (PDB-5APN) pre-60S particles were superimposed. Release of Rlp24 and Nog1-CTT would allow Rei1 to dock on its native binding site on eL22 and insert its CTT into the PET. **(c)** The Nmd3 N-terminus clashes with Nsa2 and the Nog1 N-terminal domain. Cryo-EM structures of nuclear Nog1-containing (PDB-3JCT) and late cytoplasmic Nmd3-containing (PDB-5T62) pre-60S particles were superimposed. Release of Nsa2 and the Nog1 N-terminal domain would allow 25S rRNA helix 89 to retract into its mature position and liberate the peptidyl transferase center (PTC) to enable the interaction between Nmd3-N and Tif6 and trigger Lsg1.

Strikingly, the release of another GTPase Nug2 from the pre-60S in the nucleus was reported to require its own GTPase activity, and the ATPase activity of the AAA-ATPase Rea1 (Matsuo et al., 2014). Nug2 binds the pre-60S at a site, which partially overlaps with the binding site for the NES containing adaptor Nmd3 (Sengupta et al., 2010). Only after Nug2 release, Nmd3 can be recruited to the pre-60S, a critical step that drives pre-60S nuclear export. In this case, coupled ATPase and GTPase activities, together with components of this assembly factor cluster (Figure 2), form a nuclear checkpoint to prevent a pre-60S, from prematurely acquiring export competence. Similar to Nug2, our data strongly point to Nog1 release from the pre-60S requiring its own GTPase activity and the ATPase activity of the AAA-ATPase Drg1. Failure to release Nog1 blocks a specific branch of the cytoplasmic maturation pathway critical to assemble a functional 60S subunit.

Previously, a set of multiplexed SRM-MS assays enabled the discovery of assembly factors that need to travel with the pre-60S to the cytoplasm to be released (Altvater et al., 2012). This approach involved tedious development and validation of mass spectrometric assays for every assembly factor of interest. In this study, we applied SWATH-MS to analyze proteomes of pre-ribosomal particles in an unbiased manner (Gillet et al., 2012). SWATH-MS offers similar performance characteristics as SRM-MS with the added advantage that the analysis is not limited to a set of pre-selected peptides monitored during data acquisition. The scope of quantified peptides can be extended to any peptide of interest post-acquisition due to the fact that SWATH-MS acquires fragment ion patterns for all peptides present in a sample. Using this approach, we quantified the abundance of assembly factors bound to multiple particles and organized them into functional clusters. Further, we reproduced previous findings regarding the altered composition of cytoplasmic pre-60S particles that were trapped using Drg1^DN^. This allowed the identification of assembly factors whose release or recruitment appears to be dependent on the release of the GTPase Nog1 from the pre-60S particles. These include assembly factors Sqt1, Mex67, Ecm1, Tma16 and Arb1 for which structural information within context of the pre-ribosome is lacking. Importantly, these analyses provided clues into the initial events that drive an exported pre-60S particle towards translation competence.

In contrast to their prokaryotic counterparts, eukaryotic ribosome formation requires the concerted efforts of >200 assembly factors and >40 ATPases, GTPases, ATP-dependent RNA helicases and kinases that interact with pre-ribosomal particles at distinct maturation steps. Our work flow of combining genetic trapping with SWATH-MS provides a powerful tool to uncover the intricate coordination between energy-consuming enzymes to sequentially release assembly factor clusters from maturing pre-ribosomal particles, and thereby reveal checkpoints during ribosome formation.

## Acknowledgements

VGP is supported by grants from the Swiss National Science Foundation, NCCR RNA & Disease, Novartis Foundation, Olga Mayenfisch Stiftung and a Starting Grant Award from the European Research Council (EURIBIO260676). RA is supported by ERC grant Proteomics v3.0 (AdvG233226). This work is dedicated to the memory of Dr. Cohue Peña, who unexpectedly passed away.

## Author contributions

PN and VGP designed the study. PN performed yeast functional studies with the help of MA and YC. LG and OTS performed SWATH-MS analyses in the laboratory of RA. PN, CP and VGP interpreted the results. CP, PN, LG and VGP wrote the manuscript.

## STAR Methods

### Yeast strains and plasmids

The *Saccharomyces cerevisiae* strains used in this study are listed in Table S1. Genomic disruptions, C-terminal tagging and promoter switches at genomic loci were performed according to established protocols (Janke et al., 2004; Longtine et al., 1998; Puig et al., 2001).

Plasmids used in this study are listed in Table S2. Details of plasmid construction will be provided upon request. All recombinant DNA techniques were performed according to established procedures using *E. coli* XL1 blue cells for cloning and plasmid propagation. Mutations in *NOG1* were generated using the QuikChange site-directed mutagenesis kit (Agilent Technologies, Santa Clara, CA, USA). All cloned DNA fragments and mutagenized plasmids were verified by sequencing.

### Fluorescence microscopy

For assessing the localization of PGAL1-Nog1G223A-GFP, BY4741 cells were transformed with YEP351-Nog1-GFP and YEP351gal-Nog1G223A-GFP and grown in SR medium until OD600 = 0.4-0.6, induced for 30 min with 2% galactose. Cells were then washed once in YPD and incubated in YPD for3 hours before imaging. For assessing localization of assembly factors, endogenously GFP-tagged strains were transformed with PGAL1 plasmids and were grown in SR medium until OD600=0.2-0.4 and then induced with 2% galactose for 3 hours. For Nmd3^3A^-GFP visualization, a Nmd3 shuffle strain was transformed with Nmd3^3A^ –GFP (NMD3I493A L497A L500A) and transformants were applied on FOA plates to shuffle out the wild-type *NMD3*. The resulting Nmd3^3A^-GFP strain was transformed with PGAL1 plasmids and grown the same as the endogenously GFP-tagged strains. For experiments with Diazaborine, cells were incubated 1 hour prior to imaging with either DMSO or with 370 μM Diazaborine (dissolved in DMSO; M. Peter, ETH Zürich, Switzerland) in 1 ml of cell culture for 1 hour at 30°C on shaker. When cells were ready to be harvested, the pellet was washed one with H2O. 3 μl of cells were transferred on a microscopy slide (VWR) covered with a glass slip (18×18 mm no 1, VWR).

Rpl24-TAP localization was assayed by indirect immunofluorescence. Rpl24-TAP strained were transformed with pGAL1-containing plasmids, and cultures were grown in the appropriate conditions to OD600 = 0.4-1. After adding formaldehyde (final conc. 4%) to the cultures, cells were fixed for 30 min at 30°C. Cells were then centrifuged, and incubated with 1ml of 0.1M KPi pH 6.4 with 3.7% Formaldehyde for 15 min at 30°C. Cells were then washed twice with 0.1M KPi pH 6.4 and once in spheroplasting buffer (0.1 M KPi pH7.4, 1.2 M Sorbitol, 0.5 mM MgCl2). The pellet was resuspended in 200 μl spheroplasting buffer and was either stored at -20°C or directly processed for spheroplasting. For spheroplasting, 2 μl of 1M DTT was added to the 200 μl cells and incubated for 15 min at 30°C. Zymolyase 100T was added (final conc. 50 μg/ml) to the 200 μl and incubated for 10 min. Cells were quickly checked under the microscope to see whether spheroplasts were formed (spheroplasts appear black), otherwise incubated for a longer time, but no longer than 20 min. Then, the spherolplasts were centrifuged for 2 min 2000 rpm, washed and resuspended in 100-400 μl spheroplasting buffer. At this point, spheroplasts were either stored at-20°C or directly used for antibody staining. 20 μl of poly-L-lysine (0.1% (w/v), Sigma-Aldrich) was applied per well on slides (8 well, 6 mm Menzel-Gläser Diagnostika, Brauschweig, Germany), incubated for 5 min, washed three times with dH2O, aspirated and air-dried. Approximately 40 μl of spheroplasts were added onto lysine-coated slide well. Non-adhering cell were removed after 30 seconds and the slide was incubated in an ice-cold methanol bath for 6 min before it was transferred to an ice-cold acetone bath for 10 seconds. The slide was air-dried and proceeded for blocking with 30 μl BSA/PBS (1x PBS, 1% w/v BSA) for 30 min in the dark at RT in a humid chamber. The blocking solution was removed, and primary antibody (anti-CBP 1:1000; Thermo Scientific, Rockford, IL, USA) in BSA/PBS was added to the wells and incubated for at least an hour (up to overnight) at RT in the dark. Wells were washed 3 times with BSA/PBS and incubated with secondary antibody AlexaFluor568 coupled anti-rabbit (Molecular Probes, Inc., Eugene, OR, US) at RT in the dark. After washing, cells were incubated for 30S with DAPI (1 μg/ml in BSA/PBS; Sigma-Aldrich). Wells were washed three times with BSA/PBS and dried before the slides were mounted with Mowiol (Calbiochem, San Diego, CA, USA).

Fluorescence signal was examined using a Leica DM6000B microscope fitted with a 63x 1.25 NA 1.30-0.60 NA oil immersion lens (HCX PL Fluotar; Leica). Pictures were acquired with a digital camera (ORCA-ER; Hamamatsu Photonics) and with Openlab software (Perkin Elmer). Representative sections were selected using ImageJ software and processed in AdobePhotoshop. Corresponding differential interference contrast (DIC) pictures were taken for each fluorescence image.

### Tandem affinity purifications (TAPs) and Western analyses

Whole cell extracts were prepared by alkaline lysis of yeast cells (Kemmler et al., 2009). Tandem affinity purifications (TAP) of pre-ribosomal particles were carried out as previously described (Altvater et al., 2014; Faza et al., 2012). Calmodulin-eluates were separated on NuPAGE 4-12% Bis-Tris gradient gels (Invitrogen, Carlsbad, CA, USA) and visualized by either Silver staining or Western analyses using indicated antibodies. To analyze the samples by SWATH-MS proteins TEV eluates were precipitated using trichloroacetic acid (TCA), washed once in cold acetone, air dried and processed further (see paragraphs on SWATH-MS analyses).

Western analyses were performed as previously described (Kemmler et al., 2009). The following antibodies were used: α-Mex67 (1:5,000; C Dargemont, Institut Jacques Monod, Paris, France), α-Nmd3 (1:5,000; A Johnson, University of Texas at Austin, Austin, TX, USA), α-Nog1 (1:1,000; M Fromont-Racine, Institut Pasteur, Paris, France), α-Nop7 (1:2,000; B Stillman, Cold Spring Harbor Laboratory, New York, NY, USA), α-Nug1 (1:1,000; this study), France), α-Tif6 (1:2,000; GenWay Biotech, San Diego, CA, USA), α-Yvh1 (1:4,000; Altvater et al., 2012), α-Rpl1 (1:10,000; F Lacroute, Centre de Génétique Moléculaire du CNRS, Gif-sur-Yvette, France), α-Rpl3 (1:5,000; J Warner, Albert Einstein College of Medicine, Bronx, NY, USA) and α-Rpl35 (uL29) (1:4,000; Altvater et al., 2012), α-Gsp1 (1:3,000; rabbit; Fischer et al., 2015), α-Nsa2 (1:2000; M. Fromont-Racine, Institut Pasteur, Paris, France), α-GFP (1:2000; Roche); α-FLAG (1:2000; Sigma). The secondary HRP-conjugated α-rabbit and α-mouse antibodies (Sigma-Aldrich, St. Louis, MO, USA) were used at 1:1,000-1:5,000 dilutions. Protein signals were visualized using Immun-Star HRP chemiluminescence kit (Bio-Rad Laboratories, Hercules, CA, USA) and captured by Fuji Super RX X-ray films (Fujifilm, Tokyo, Japan).

## SWATH-MS analyses

### Sample preparation for the SWATH-MS analyses

The proteins were solubilized in a denaturing buffer containing 8 M urea and 0.1 M NH4HCO3. They were then reduced with 12 mM DTT at 37°C for 30min and alkylated with 40 mM iodacetamide at room temperature in the dark for 30min. The samples were then diluted with 0.1 M NH4HCO3 to reach a final urea concentration of 1M and digested with sequencing grade porcine trypsin (Promega, 1:100 trypsin:protein). Digestion was stopped by adding formic acid to a final concentration of 1% (pH ~2). Peptides were desalted using macro spin columns (Nest group) according to the following procedure: Cartridges were wetted with one volume (350 μl) 100% methanol, washed with two volumes of 80% acetonitrile, 0.1% formic acid (FA) and equilibrated with three volumes of 0.1% FA. The acidified peptides were loaded twice on the cartridge, washed with three volumes 0.1% FA and eluted with two volumes 50% acetonitrile, 0.1% FA. Peptide were dried in a speedvac concentrator and resolubilized in 10 μl of 0.1% formic acid. The samples were then transferred to an MS vial and spiked with 1:20 (v:v) of iRT peptides (Escher et al., 2012).

### Mass spectrometry data acquisition

1 μg of peptides were injected on a 5600 TripleTof mass spectrometer (ABSciex, Concord, Ontario) interfaced with an Eksigent NanoLC Ultra 1D Plus system (Eksigent, Dublin, CA). The peptides were separated on a 75-μm-diameter, 20 cm long New Objective emitter packed with Magic C18 AQ 3 μm resin (Michrom BioResources) and eluted at 300nl/min with a linear gradient of 5-to-35% Buffer A for 120min (Buffer A: 2% acetonitrile, 0.1% formic acid; Buffer B: 98% acetonitrile, 0.1% formic acid). MS data acquisition was performed in either data-dependent acquisition (DDA, top20, with 20 s dynamic exclusion) or data-independent acquisition (DIA) SWATH-MS mode (32 fixed precursor isolation windows of 25Da width (+1 Da overlap) each acquired for 100ms plus one MS1 scan acquired for 250ms) as described in (Gillet et al., 2012). The mass ranges recorded were 360-1460 m/z for MS1 and 50-2000 m/z for MS2. For either mode, the collision energy was set to 0.0625 × m/z -6.5 with a 15-eV collision energy spread regardless of the precursor charge state.

### SWATH-MS assay library generation

The DDA data recorded as described above were used to generate an assay library essentially as described (Schubert et al., 2015). In short, the raw DDA files were converted to mzXML using the qtofpeakpicker component of msconvert (ProteoWizard v 3.0.9987). The converted files searched with Comet (2014.02 rev. 0) and Mascot (version 2.5.1) using the yeast SGD database (release 13.01.2015) containing 6,713 proteins plus one protein entry for the concatenated sequence of the iRT peptides and as many decoy protein entries generated by pseudo-reversing the tryptic peptide sequences. The search parameters were as follows: +/−25ppm tolerance for MS1 and MS2, fixed cysteine carbamidomethylation, variable methionine oxidation, semi-tryptic and up to 2 missed cleavages per peptide allowed. The Comet and Mascot search results were further processed using peptideProphet (Keller et al., 2002) and aggregated using iProphet (Shteynberg et al., 2011) (TPP v4.7 rev 0). The search results were filtered for an iProphet cutoff of 0.877603, corresponding to a 1% protein false discovery rate (FDR) estimated by MAYU (Reiter et al., 2009). The search results contained 22,961 peptides matching to a set of 2,610 proteins. The consensus spectral library was generated using SpectraST (Lam et al., 2008) and the assay library thereof was exported using the spectrast2tsv.py script (Schubert et al., 2015) with the following parameters: 6 highest intensity fragments (of charge 1+ or 2+) per peptide, within the mass range 350-2000 m/z and excluding the fragments within the precursor isolation window of the corresponding swath. The final library contains assays for 23,960 peptide precursors (thereof 21,472 proteotypic precursors covering 2,209 unique proteins). The assay library was exported to TraML format with shuffled decoys appended as described (Schubert et al., 2015).

### SWATH-MS data analysis

The SWATH-MS data was extracted with the above-mentioned assay library through the iPortal interface with openSWATH (Rost et al., 2014) (openMS 2.1.0), pyProphet (Teleman et al., 2015) and TRIC alignment (Rost et al., 2016) using the same parameters as described in (Navarro et al., 2016). The SWATH identification results were further filtered to keep all the peptide assays with m-score below 0.01 for the protein entries with at least one peptide with an m-score below 0.00000208508 (corresponding to a protein FDR of 1%). A set of ribosomal proteins (RPL28, RPL14B, RPL14A, RPL3, RPL19A/RPL19B, RPL35A/RPL35B, RPL12A/RPL12B, RPL27A/RPL27B, RPL2B/RPL2A, RPL14A/RPL14B, RPL7B/RPL7A, RPL1A/RPL1B) was used to normalize (mean-centering) the data as described in (Altvater et al., 2012). Only the proteotypic peptides were then kept, as well as the assays identified in the three triplicates for at least one AP-MS condition. Finally, proteins with less than 2 peptides were filtered out. The missing values for each peptide assay were imputed using a random value between 0.7 and 0.9 fold the lowest intensity of that peptide assay throughout the dataset. All the peptide assays values were then summed to a protein intensity value and a protein intensity mean per AP-MS condition. For the general protein heatmaps, each protein intensity was normalized to that of the highest protein value for the reported set. For the bar plots, the fold changes were calculated for each protein to that of the AP-MS condition with the highest protein intensity while the standard deviations were calculated per condition.

### Data availability

SWATH-MS data have been deposited to the ProteomXchange Consortium via the PRIDE partner repository with the dataset identifier PXD011382.

